# Spatial reconstruction of single enterocytes uncovers broad zonation along the intestinal villus axis

**DOI:** 10.1101/261529

**Authors:** Andreas E. Moor, Yotam Harnik, Shani Ben-Moshe, Efi E. Massasa, Keren Bahar Halpern, Shalev Itzkovitz

## Abstract

The intestinal epithelium is a highly structured tissue composed of repeating crypt-villus units^1,2^. Enterocytes, which constitute the most abundant cell type, perform the diverse tasks of absorbing a wide range of nutrients while protecting the body from the harsh bacterial-rich environment. It is unknown if these tasks are equally performed by all enterocytes or whether they are spatially zonated along the villus axis^3^. Here, we performed whole-transcriptome measurements of laser-capture-microdissected villus segments to extract a large panel of landmark genes, expressed in a zonated manner. We used these genes to localize single sequenced enterocytes along the villus axis, thus reconstructing a global spatial expression map. We found that most enterocyte genes were zonated. Enterocytes at villi bottoms expressed an anti-bacterial Reg gene program in a microbiome-dependent manner, potentially reducing the crypt pathogen exposure. Translation, splicing and respiration genes steadily decreased in expression towards the villi tops, whereas distinct mid-top villus zones sub-specialized in the absorption of carbohydrates, peptides and fat. Enterocytes at the villi tips exhibited a unique gene-expression signature consisting of Klf4, Egfr, Neat1, Malat1, cell adhesion and purine metabolism genes. Our study exposes broad spatial heterogeneity of enterocytes, which could be important for achieving their diverse tasks.

## Main Text

The intestinal tract is responsible for nutrient digestion and absorption, secretion of mucus and hormones, interactions with commensal microbiota and protection of the organism from pathogenic microbes^1,2^. This wide array of tasks requires the presence of different cell types that are specialized for their respective functions. Enterocytes, which represent the majority of cells in the epithelial layer, constantly migrate along the villi walls until they are shed off from their tips 3-5 days after their emergence from crypts. The positions of enterocytes along the villus axis correlate with their age^3^, exposure to morphogen gradients^1^ and hypoxia^4^, yet the positional effects on enterocyte function are largely unknown. Previous work investigated transcriptomic changes along the small intestinal crypt-villus axis with bulk samples and DNA microarray-based expression profiles in mouse^5,6^ and human tissue^7^. This body of work revealed some broad compositional differences of the crypt and the villus, yet its low spatial resolution (comparing bulk crypts to bulk villi), uncontrolled mixes of different cell types and the low sensitivity of microarray-based transcriptomics precluded the detection of spatial expression changes and heterogeneity of enterocytes along the villus.

Single-cell RNA sequencing (scRNAseq) has revolutionized our ability to characterize individual cells in-depth^8^; it was recently utilized in the intestine to identify cell types^9^ and sub-populations of intestinal stem cells^10^, tuft cells^11,12^ and enteroendocrine cells^9,12–14^. However, spatial heterogeneity and specialization along the villus axis within the enterocytes, the largest cell compartment, has not been addressed. Relating such heterogeneity to tissue coordinate is challenging, as the spatial origin of individual cells is lost when the tissue is dissociated for scRNAseq. We and others have developed approaches to spatially reconstruct scRNAseq data by making use of known expression profiles of landmark genes characterized by RNA in-situ hybridization^15–18^ (Fig. 1a). This approach is infeasible, however, when no prior knowledge exists regarding zonated landmark genes. Here we established a comprehensive panel of landmark genes, characterized by RNAseq of laser-capture-microdissected epithelial samples originating from differential villus zones. We used these to reconstruct the spatial tissue coordinates of enterocytes in scRNAseq data and uncovered vast heterogeneity and sub-specialization along the villus axis. This spatial division of labor of mature enterocytes could be optimal for achieving the diverse tasks of this tissue^3^.

**Figure 1:**
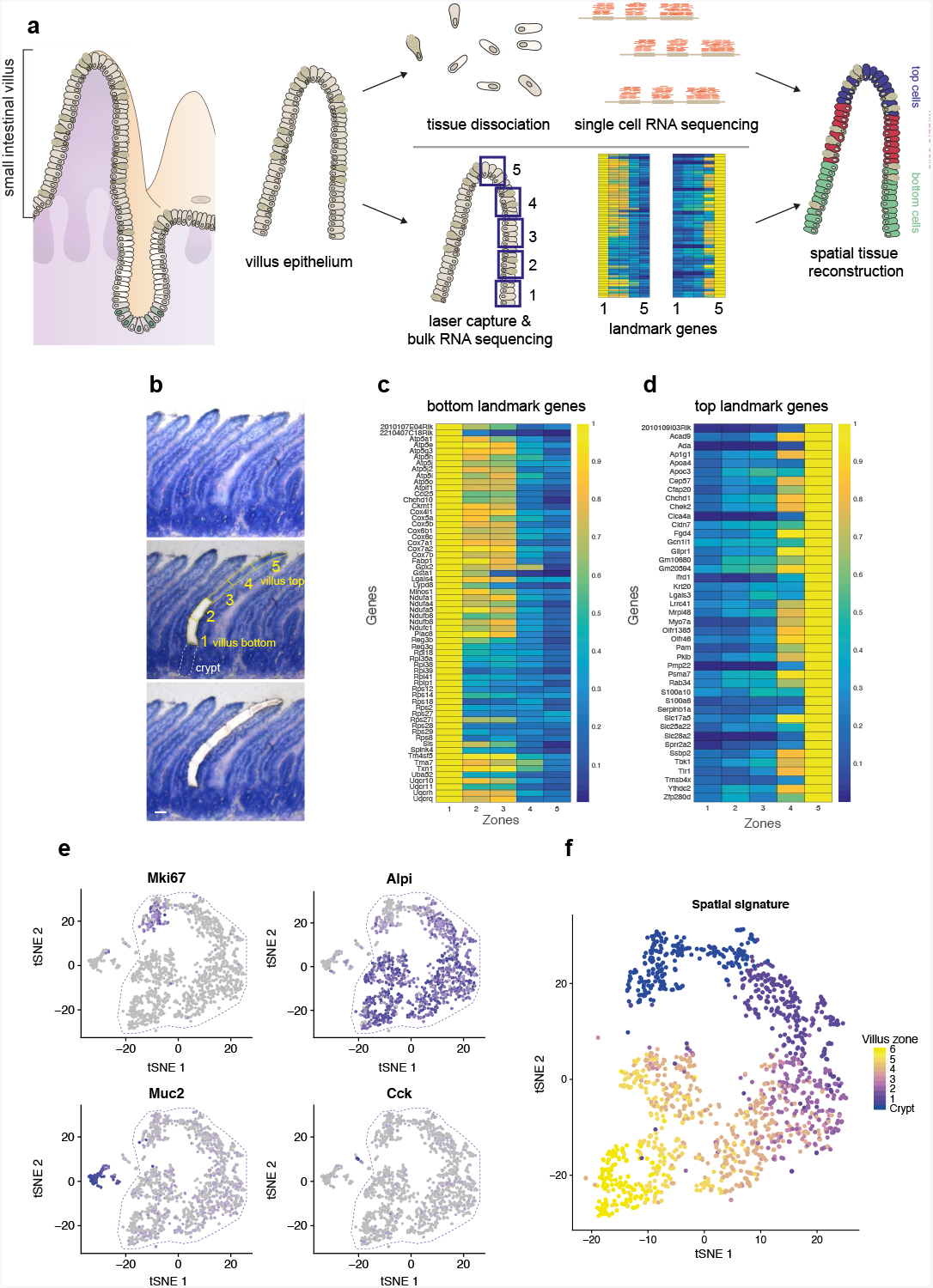
Spatial reconstruction of villus enterocytes. **a**,Scheme of the experimental approach. The villus epithelium is dissociated into single cells; these cells are profiled by scRNAseq. In parallel, spatial landmark genes are retrieved by bulk RNAseq of villus quintiles obtained using laser capture microdissection (LCM). The original position of the sequenced single cells is then inferred based on their expression levels of the landmark genes. **b**, Laser capture microdissection of villus epithelium quintiles. Scale bar 50μm. **c**, LCM-RNAseq expression of the villus-bottom landmark genes. **d**, LCM-RNAseq expression of the villus-top landmark genes. Profiles in c-d are normalized to the maximum expression level for each gene. **e**, tSNE plots of intestinal marker gene expression in single Lgr5-eGFP negative cells^13^. Mki67 is expressed in transient amplifying cells in the crypt, Alpi in villus enterocytes, Muc2 in goblet cells and Cck in enteroendocrine cells. Depicted analyses are based on raw data from NCBI GEO datasets GSM2644349 and GSM264435013. **f**, tSNE plot of the enterocyte and progenitor populations in the sorted Lgr5 negative dataset. Each cell is colored according to its inferred villus zone.

To extract a panel of enterocyte landmark genes, we used laser capture microdissection (LCM) to isolate epithelial cells from five equally spaced compartments between the bottom and tops of villi in the mouse Jejunum (Fig. 1a,b). Bulk RNA sequencing (RNAseq) of these isolated villus quintiles revealed genes with decreasing (Fig. 1c) and increasing (Fig. 1d) expression gradients. We defined a set of 62 villus-bottom landmark genes and 43 villus-top landmark genes to be used for spatial reconstruction of scRNAseq data (Supp. Fig. 1).

We used our LCM-RNAseq reference to identify a scRNAseq dataset^13^, which included enterocytes that spanned the entire villus axis (Supp. Fig. 2). Mature and progenitor enterocytes were clearly demarcated by the expression of Alpi and Mki67 respectively (Fig. 1e). We assigned each sequenced mature enterocyte a unit-less spatial coordinate x that was based on the ratio between the summed expression of the top and bottom landmark genes. For each cell, x correlated with its position along the villus axis (Methods, Supp. Fig. 2). By computing the x values of the five laser captured areas we were able to assign each cell to one of 6 zones from the bottom to the top of the villus (Fig. 1f). We averaged, for every gene, the expression of single cells in each of these zones, to obtain a comprehensive spatial map of gene expression along the intestinal villus (Fig. 2a,b).

**Figure 2:**
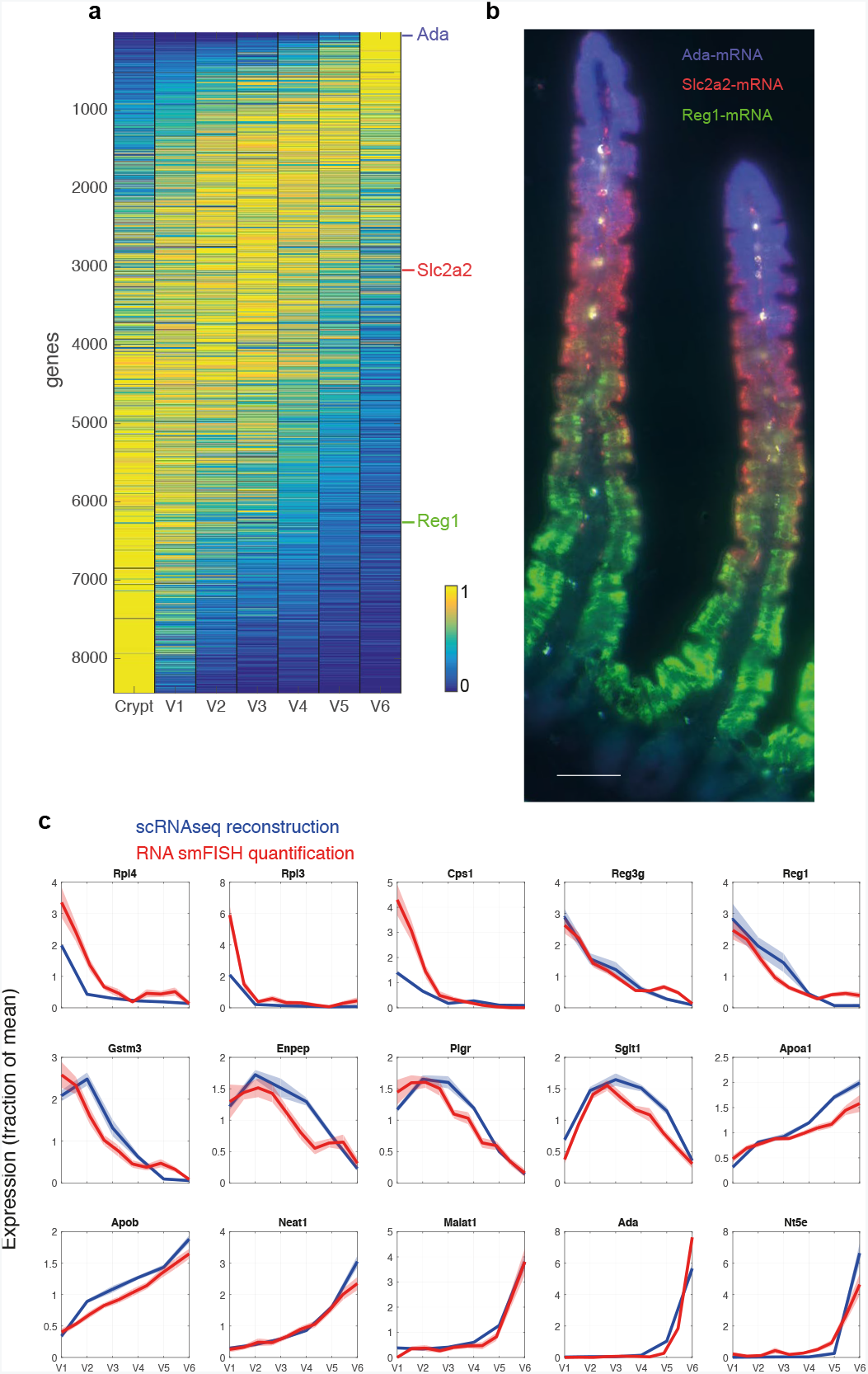
A global zonation map of enterocytes along the villus axis. **a**,Spatially-reconstructed gene expression zonation profiles. Profiles are normalized to their maximum and sorted according to their center of mass. Shown are the genes with maximal zonation value above 10^−5^. **b**, smFISH staining of Ada mRNA (blue) that is expressed in villus tip enterocytes, Slc2a2 mRNA (red) that is expressed in villus middle enterocytes and Reg1 mRNA (green) that is expressed in villus bottom enterocytes. Scale bar 50μm. **c**, Validation of the reconstructed zonation profiles using smFISH. Dark blue line depicts scRNAseq mean expression level, light blue area denotes its SEM (standard error of the mean). Dark red line depicts smFISH mean expression level, light red area denotes its SEM. All profiles are normalized by their means across zones. SmFISH profiles based on measurements from at least 10 villi from 3 different mice.

Our spatial reconstruction included more than 9,832 enterocyte expressed genes, 8,126 of which (83%) were significantly zonated (Methods, q-value < 0.05). Thus, differentiated enterocytes exhibit ubiquitous spatial heterogeneity with only a small minority of genes invariably expressed from the bottom to the top of the villi. We used single molecule Fluorescence in-situ Hybridzation^17^ (smFISH) to validate our predicted zonated expression profiles for 15 enterocyte genes, demonstrating the accuracy of reconstruction (Fig. 2c, Supp. Fig. 3-5,7,8).

We used k-means to cluster the genes into five distinct groups, ordered from villus bottom to top according to their average zonation profiles (Fig. 3a). Gene set enrichment analysis^19^ revealed enriched gene ontology (GO) terms for each cluster (Fig. 3a). Cluster 1 contained genes that decreased progressively from Crypts to villi tips. These included a global decline of translation, transcription and RNA splicing genes. Thus, enterocyte biosynthetic capacity seems to be gradually decreasing as enterocytes migrate along the villus axis. Mitochondrial GO terms were enriched in cluster 2 (Fig. 3a, b). This decrease in mitochondrial content may be an adaptation to the decreasing gradient of oxygen concentration, previously demonstrated along the villus axis^4^. Cluster 2 also contained glutathione transferase activity, which contains Gstm3 (Fig. 2c, Fig. 3a, Supp. Fig. 5), as well as acute phase response genes such as Reg genes, which were highly expressed at the villus bottom yet barely expressed in the adjacent crypt (Fig. 3c). Cluster 3 consisted of intestinal transport annotations, which peaked at the mid-villus zones. Genes in cluster 4 increased in expression up to the mid-villi zones and included many brush border components. Cluster 5 included lipoprotein biosynthesis and cell adhesion processes as well as the lncRNA markers of speckles (Malat1, Supp. Fig. 4) and paraspeckles (Neat1, Supp. Fig. 4), all monotonically increasing towards the villi tips.

**Figure 3:**
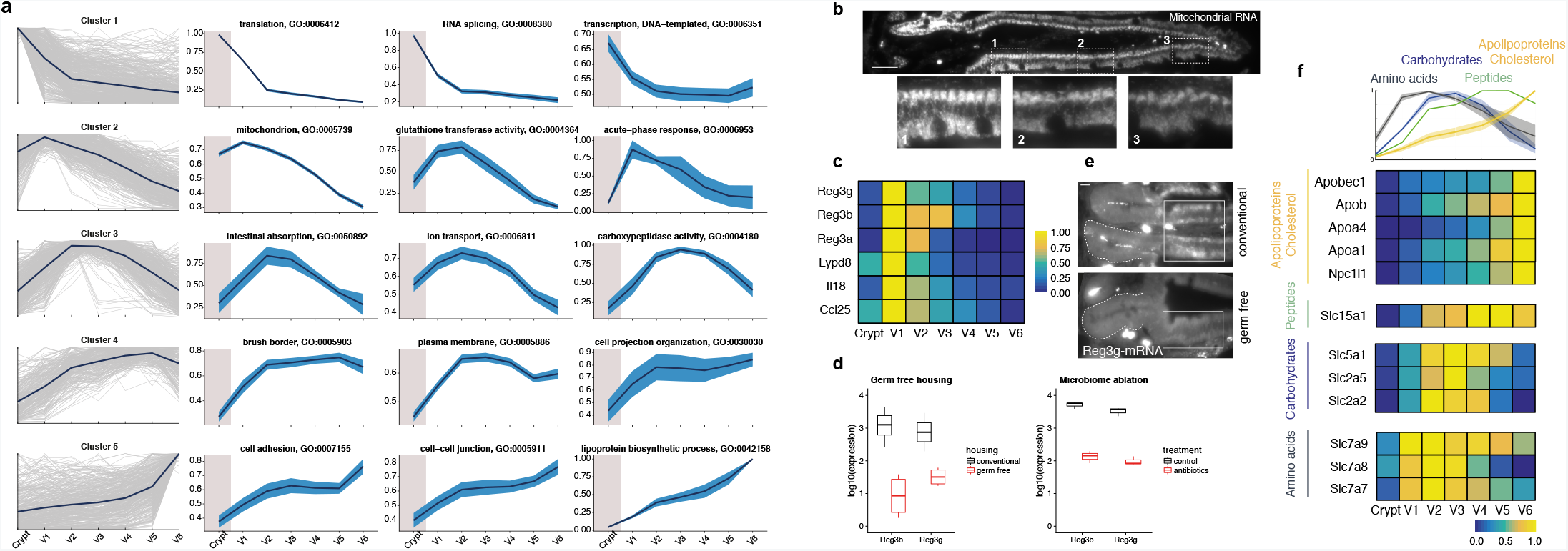
Functional sub-specialization of villus enterocytes. **a**,Left column: 5 clusters of genes with similar zonation profiles along the villus length. Blue line represents cluster mean; grey lines depict individual genes. Columns 2 to 4: representative gene ontology (GO) terms enriched in the identified gene clusters. Dark blue line show GO term mean; light blue area denotes SEM. **b**, smFISH of mitochondrial light-strand RNA. Enlarged inserts show the gradual decrease in mitochondrial light-strand RNA expression from villus bottom to villus top. Scale bar 50μm. **c**, Heatmap of zonation profiles of Reg gene family members as well as other peptides involved in microbiota-host interactions. **d**, Left: RNAseq expression levels of Reg3b and Reg3g of mice that are in conventional or germ free environments. Analysis is based on raw data from the NCBI GEO dataset GSE8112524. Right: RNAseq expression levels Reg3b and Reg3g of mice that were treated with vehicle or with antibiotics to ablate the microbiome. Analysis is based on raw data from the NCBI GEO dataset GSE7415725. Horizontal lines are medians, boxes are 25-75 percentiles and vertical lines denote values within 1.5 x interquantile range of the respective quartile. **e**, smFISH of Reg3g expression in the crypt and adjacent bottom villus area in mice that were either kept conventionally or in germ free conditions. Dashed line denotes the crypt, solid box indicates the villus bottom zone where Reg3g is expressed. Scale bar 10μm. **f**, Heatmaps and averaged zonation profiles of selected genes that are important for the absorption of distinct nutrient classes. Black line: amino acid transporters, Blue line: carbohydrate transporters, green line: the peptide transporter Slc15a1, yellow line: fat and cholesterol absorption. Profiles are normalized to their maximal value across zones.

Genes of the Reg family belong to the calcium-dependent lectin genes and encode small secretory proteins. Reg3 proteins bind preferentially to Gram-positive bacteria and are bactericidal^12,20^. Our spatial reconstruction uncovered a restricted zone at the bottom of the villus in which enterocytes strongly expressed Reg family members, as well as other peptides involved in microbiota-host interactions such as Lypd8^21^, Ccl25^22^ and Il18^23^ (Fig. 3c). Reg3b and Reg3g are the two most significantly downregulated genes when comparing RNAseq of conventional to germ free mice^24^ (Fig. 3d, Supp. Fig. 6). Their expression is strongly decreased upon microbiome ablation with antibiotic intervention^25^ (Fig. 3d). We used sm-FISH to demonstrate a sharp decrease in Reg3g at the bottom of the villus in germ free mice compared to controls (Fig. 3e, Supp. Fig. 6b). These findings reveal an antimicrobial zone above the crypt that might function as a gatekeeper for the crypt stem-cell niche to minimize its exposure to pathogenic microbes.

Enterocytes absorb a wide range of nutrients, including carbohydrates, amino acids and lipids. We found that the transporters for these key nutrient families were non-overlapping in their expression domains. Whereas amino acid and carbohydrate transporters were enriched at the middle of the villus (Fig. 3a,f), Slc15a1 that encodes the main peptide transporter Pept1 was shifted in expression towards the upper villus zones, and the cholesterol transporter Npc1l1 and the lipoprotein biosynthesis machinery, necessary for the assembly of chylomicrons peaked in expression at the villi tips (Fig. 3f). The zonated expression of lipoprotein genes at the villi tops can explain previous findings of higher chylomicron density at the villi tips shortly after lipid gavage^26^. Thus enterocytes seem to be sub-specialized in preferential nutrient absorption according to their position along the villus axis.

Our spatial reconstruction revealed a sharp increase in the expression of distinct signaling pathways and transcription factor sets at the villi tips. These genes included Egfr, Klf4 and the AP-1 transcription factors Fos and Junb (Fig. 4a,b). Egfr signaling has been implicated in tight junction organization^27^. Its increased expression at the villi tips might initiate reorganization of cell adhesion (Fig. 3a, Supp. Fig. 7) that is necessary for subsequent cellular shedding. Villus tip cells further expressed a signature of purine catabolism genes, including Enpp3, Nt5e, Slc28a2 and Ada (Fig. 4c-f, Supp. Fig. 8). Enpp3 and Nt5e, which are encoding ecto-nucleotidases that convert ATP to AMP and AMP to adenosine^28^ respectively were expressed in a sequential manner at the villi tips. Enpp3 increased steeply from villi zones 4 to 6, whereas Nt5e was only expressed in zone 6 at the very tips of the villi. (Fig. 4f). The tip-enriched gene Slc28a2 encodes a Na+ coupled high affinity adenosine transporter^29^ that could shuttle the generated adenosine into the cytosol. There, adenosine can be converted to inosine by the adenosine deaminase (Ada), which we also found to be confined in expression to this zone (Fig. 4d). We validated the tip-enriched expression of Nt5e protein (also known as Cd73) and observed that most of this ecto-nucleotidase was localized to the luminal side of villus tip enterocytes (Fig. 4e). Bacterially dependent luminal ATP is a danger signal that activates intestinal immune cells^30^, whereas adenosine and inosine, the products of the revealed villus tip signaling program, exert potent anti-inflammatory functions in the intestine^31^. Thus the villus tip expression program could be important for maintaining a balanced immune response to the microbiome.

**Figure 4:**
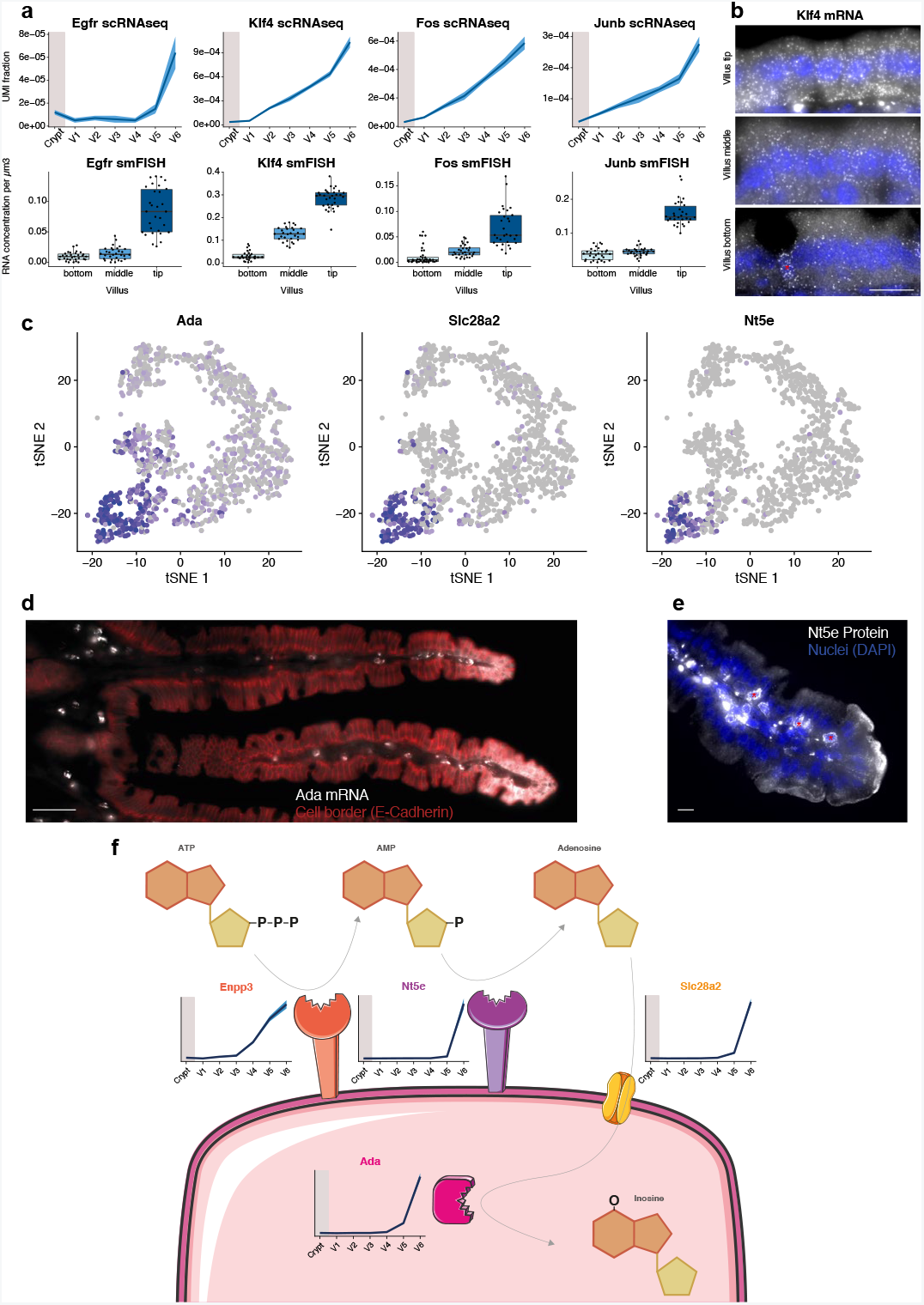
Spatial reconstruction reveals a villus tip expression program. **a**,Signaling and transcriptional programs of the villus tip zone. Top row: scRNA-seq-inferred expression profiles. Dark blue line: mean expression, light blue area: SEM. Bottom row: quantification of smFISH expression in bottom, middle and top part of the villus. **b**, smFISH of Klf4 mRNA expression in the bottom, middle and top part of the villus. Asterisk in the villus bottom field of view marks a goblet cell (known to express Klf4). Scale bar 10μm. **c**, tSNE plots of the expression patterns of Ada, Slc28a2 and Nt5e, three of the identified purine catabolism villus tip marker genes. **d**, smFISH of Ada mRNA (white), cell borders (E-Cadherin protein) are depicted in red. Scale bar 50 μm. **e**, immunofluorescence staining of Nt5e protein at the villus tip, demonstrating apical localization of the protein. Asterisks mark extracellular Nt5e proteins on intraepithelial lymphocytes. Scale bar 10 μm. **f**, model of functional interaction of villus tip genes in purine catabolism. Luminal ATP is hydrolyzed to AMP by Enpp3 and subsequently converted to Adenosine by Nt5e. Part of this generated Adenosine can be absorbed by the high-affinity Adenosine transporter Slc28a2. Intra-cellular Ada converts Adenosine to Inosine.

Our study uncovered an unexpectedly broad spatial heterogeneity within small intestinal enterocytes. The large majority of genes are significantly zonated along the villus-axis. The secluded stem cell niche in the intestinal crypt seems to be protected by a layer of gatekeeper enterocytes that secrete antibacterial Reg proteins at its entrance. These enterocytes may complement the secretory Paneth cells in the protection of the crypt resident stem cells. The absorption machinery of specific nutrients is compartmentalized in distinct villi zones, potentially leading to more efficient nutrient uptake. Villus tip cells appear to orchestrate an immune-modulatory program that might have important implications for host-microbe interactions in health and disease.

Our spatial reconstruction revealed a wide array of surface markers such as Cd73 (Supp. Fig. 9), that can be used in flow cytometry experiments to isolate bulk, spatially-stratified enterocyte populations. This could enable deeper characterization of the epigenome, proteome, metabolome and mutation signatures along the villus spatial axis. The broad heterogeneity we observed along the villus axis seems to be conserved among the three small intestinal regions (Supp. Figure 10). It would be interesting to use our approach to explore the zonation profiles of enterocytes in diverse intestinal pathologies. More generally, the use of LCM-RNAseq to extract a large set of landmark genes in an unbiased manner, is a generic alternative to FISH-based single-cell spatial reconstructions^15–18^, particularly useful when no prior knowledge on zonation exists. This could be used to reconstruct expression cell atlases of other tissues and tumors^32,33^.

## Materials and Methods

### Animal experiments

All mouse experiments were conducted in accordance with institutional guidelines and approved by the Institutional Animal Care and Use Committee of the Weizmann Institute of Science, Rehovot. Experiments were performed with 8-12 week old male C57BL/6 mice that were obtained from the Harlan laboratories or the WIS animal breeding center and were housed in individually ventilated cages. Mice were fed regular chow ad libitum and were exposed to phase-reversed circadian cycles. Germ-free C57BL/6 mice were housed in sterile isolators^34^. Mice were sacrificed and the proximal Jejunum was flushed with cold PBS, laterally cut, spread on dry whatman filter paper with the villi facing upwards and cut into rectangles with a length of 1.5cm. Flat tissue on whatman paper was fixed in 4% paraformaldehyde (PFA) in PBS for 3 hours and subsequently agitated in 30% sucrose, 4% PFA in PBS overnight at 4°C. Fixed tissues were embedded in OCT (Tissue-Tek). We found that flat embedding of Jejunum pieces was important for preserving the intact morphology of full-length villi.

### smFISH

8um thick sections of fixed proximal Jejunum were sectioned onto poly L-lysine coated coverslips and used for smFISH staining. Probe libraries were designed using the Stellaris FISH Probe Designer Software (Biosearch Technologies, Inc., Petaluma, CA), see Supp. Table 7. The intestinal sections were hybridized with smFISH probe sets according to a previously published protocol^35^. DAPI (Sigma-Aldrich, D9542) and a FITC-conjugated antibody against E-Cadherin (BD Biosciences, 612131) were used as nuclear and cell-membrane counterstains, respectively. SmFISH imaging was performed on a Nikon-Ti-E inverted fluorescence microscope with 60x or 100× oil-immersion objectives and a Photometrics Pixis 1024 CCD camera using MetaMorph software as previously reported^35^.

Probe libraries for messenger RNAs of interest were coupled to Cy5 and Alexa594, full-length villi were identified by the presence of co-stained villus tip maker gene expression (Nt5e or Ada coupled to TMR) on the same section. smFISH signal detection requires 60x or 100x magnifications, hence several fields of view were stitched together to cover the whole crypt-villus unit. Stitching was performed with the fusion mode linear blending and default settings of the pairwise stitching plugin^36^ in Fiji^37^. Stitched villi were cropped rectangularly and underlaid with black background (which is visible in the stitched images in areas that lack data).

We used two different methods to quantify the expression profiles of transcripts along the villus-axis from the smFISH images, depending on the abundance of the transcripts of interest. For low abundance genes, dots were counted using custom Matlab program^38^ (MATLAB Release 2016a, The MathWorks Inc., USA). The bottom, top and lateral epithelial borders of each quantified villus were manually segmented based on nuclear and cell-membrane counterstains. The epithelium was automatically further segmented into 20 units from bottom to top of the villus and mRNA density (number of mRNA per unit volume, for low abundance genes) or mRNA signal intensity (mean background-subtracted intensity in segmented unit, for high abundance genes) was computed along the villus-axis. For each transcript, we quantified at least 10 villi from 3 different mice.

### Immunofluorescence

8um thick sections of fixed proximal Jejunum were sectioned onto poly L-lysine coated coverslips and fixed with cold methanol for 20 minutes. Sections were briefly washed 3 times with PBST (1xPBS, 1% BSA and 0.1% Tween 20) and were further incubated 10 minutes in PBSTX (1X PBS, 0.25% Triton 100X and 1% BSA) at room temperature for permeabilization. After 3 PBST washes, sections were blocked with PBS supplemented with 0.1% Tween 20 and 5% Normal Horse Serum for 1h at room temperature, followed by an overnight incubation at 4°C with Alexa Fluor 647 rat anti mouse CD73 conjugated antibody (BD biosciences, 561543), 1:50. Sections were then washed again with PBST 3 times and were incubated with DAPI (1:200 in PBS) for 20 minutes. Prior to mounting on slides, sections were washed with glox buffer (10mM Tris pH 8, 2x SSC, 0.4% glucose). Imaging was carried out using the same setting as for the smFISH experiments.

### Laser capture microdissection

Tissue blocks for microdissection were obtained from three 8 week-old male C57BL/6 mice. The proximal Jejunum was briefly washed in cold PBS and embedded and frozen in OCT without fixation. 8μm thick sections were cut from the frozen block, mounted on polyethylene-naphthalate membrane-coated glass slides (Zeiss, A4151909081000), air-dried for 1m at room temperature, washed in 70% ethanol (30s), incubated in water (Sigma-Aldrich, W4502, 30s), stained with HistoGene Staining Solution (ThermoFisher Scientific, KIT0401, 20s), washed vigorously in water for a total of 30s. The stained sections were dehydrated with subsequent 30s incubations in 70%,95% and 100% ethanol and air dried for 3m before microdissection.

Tissue sections were microdissected on a UV laser-based PALM Microbeam (Zeiss). The system makes use of a pulsed UV laser that cuts the tissue at indicated marks with minimal damage to surrounding cells; the cutting was performed with the following parameters: PALM 20X lens, cut energy 48 (1-100), cut focus 65 (1-100). Tissue fragments were catapulted and collected in 0.2ml adhesive cap tubes (Zeiss, A4151909191000) with these settings: LPC energy 67 (1-100), LPC focus 67 (1-100). The capturing success was visually confirmed by focusing the PALM on the targeted adhesive cap after the collection session. 8-10 Villi above 500μm length selected for microdissection for each of three mice, their villus epithelium was divided into 5 segments of equal length and isolated. A total of 30’000-45’000 μm2 of villus epithelium area was collected for each of the five villus zones per mouse.

### RNA sequencing

Library preparation for microdissected tissues was performed based on a previously published protocol^39^ with minor modifications. Specifically, we resuspended microdissected fragments in 9.5μl H2O and 1μl of 10x reaction buffer of the SMART-Seq v4 Ultra Low Input RNA Kit (Clontech, 634888) in the adhesive cap of the collection tubes. Tissue lysis was achieved by incubation for 5m at room temperature; the lysed samples were flash frozen until library preparation. The RNA was amplified with the SMART-Seq v4 kit according to the manufacturer’s instructions and by using 15 PCR cycles for the cDNA amplification step. 1ng of the amplified cDNA was converted into sequencing library with the Nextera XT DNA Library kit (Illumina, FC-131-1024). The quality control of the resulting libraries was performed with an Agilent High Sensitivity D1000 ScreenTape System (Agilent, 5067-5584). Libraries that passed quality control were loaded with a concentration of 1.8pM on 75 cycle high output flow cells (Illumina, FC-404-2005) and sequenced on a NextSeq 500 (Illumina) with the following cycle distribution: 8bp index 1, 8bp index 2, 38bp read 1, 38bp read 2.

### Bulk RNA sequencing analysis

Illumina output files were demultiplexed with bcl2fastq 2.17 (Illumina) and the resulting fastq files of mRNA sequencing experiments were pseudoaligned with Kallisto 0.43.0^40^ to a transcriptome index of the GRCm38 release 90 (Ensembl), filtered to transcripts with a source entry of “ensembl_havana”. The following flag was used for kallisto: -b 100. Sleuth 0.28.1^41^ running on R 3.3.2 was utilized to create a TPM table (Transcripts Per Million) for each sample, according to the Kallisto pseudoalignments (Supp. Table 1).

### Single cell RNA sequencing (scRNAseq) analysis

Two Lgr5-eGFP negative scRNAseq (Chromium, 10x Genomics) datasets were acquired from the NCBI GEO dataset browser (accessions GSM2644349 and GSM264435013). scRNAseq analysis was performed with Seurat 2.1.0^15^ in R 3.3.2. Cells were filtered based on mitochondrial gene content, unique molecular identifier (UMI) counts were log-normalized according to default Seurat settings. Variable genes were identified (FindVariableGenes, parameters: x.low.cutoff = 0.0125, x.high.cutoff = 3, y.cutoff = 0.5) and the following three sources of variation were regressed out: UMI number, biological replicate number and mitochondrial gene content. Principle Component Analysis was performed on the expression levels of the detected variable genes. The first 10 principal components were included for further downstream analyses based on visual inspection of Seurat’s PCElbowPlot. Cells were clustered based on the principal component analysis with the following granularity parameters: dims.use = 1:10, resolution = 1.3. Non-linear dimensional reduction (tSNE) was used to visualize the previously computed clusters. Mature enterocytes and transient amplifying clusters were identified based on Alpi and Mki67 expression, respectively. A few mis-assigned goblet, tuft and Paneth cells were removed by filtering based on expression of the following marker genes: Muc2 and Hepacam2 (Goblet), Dclk1 (Tuft) and Lyz1 (Paneth). Raw UMI counts of the resulting 1383 enterocytes and transient amplifying cells were exported and utilized for zonation reconstruction algorithm (Supp. Tables 2 and 3).

### Zonation reconstruction algorithm

To reconstruct the zonation profiles from the scRNAseq data we used the summed expression of the landmark gene (LM) panels to infer the locations of each sequenced enterocyte along the villus spatial axis. Each cell i was assigned a spatial coordinate 0≤x_i_≤1, which correlated with its location along the villus axis and was computed as the ratio of the summed expression of the top landmark genes (tLM), and the summed expression of the bottom (bLM) and top LM genes to yield

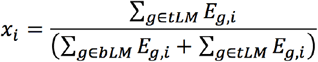

Where E_g,i_ is the expression of gene g in cell i in units of fraction of total cellular UMIs. To map xi values to spatial coordinates along the villus axis, we used the same equation to calculate the coordinate x_LCMi_ for each of the five Laser-capture-microdissected villus zones LCMi. We assigned each cell to one of 6 villus zones, V1..V6 as follows: cells with x_i_<x_LCM1_ were assigned to V1, cells with x_LCM1_≤x_i_<x_LCM2_ were assigned to V2, cells with xL_CM2_≤x_i_<x_LCM3_ were assigned to V3, cells with x_LCM3_≤x_i_<x_LCM4_ were assigned to V4, cells with x_LCM4_≤x_i_<x_LCM5_ were assigned to V5 and cells with x_LCM5_≤x_i_ were assigned to V6. For each gene and zone we calculated the means and standard errors of the means (SEM) of the expression of all genes over the cells assigned to the respective zone. Crypt gene expression was computed by the mean and SEM over the expression of single cells assigned by Seurat to the two transient amplifying clusters (Supp. Table 4).

To compute zonation significance, we used the profile’s dynamic range, defined as the difference between the maximum and minimum values of the mean-normalized profile along V1-V6 as a summary statistic. We compared the dynamic range to those obtained for 1,000 datasets in which the cells’ assigned zones were randomly reshuffled. We included genes with maximal zonation larger than 5^*^10^−6^ when computing the fraction of zonated genes. For each gene, we calculated Z-scores for the dynamic range and used the normal distribution to obtain p-values. We used Storey’s method to compute q-values (Supp. Table 4).

### Comparison of different intestinal regions

Comparison of zonation profiles among the three small intestinal regions (Supp. Fig. 10) was performed on scRNAseq data from Haber et al.^12^ (https://portals.broadinstitute.org/single_cell/data/public/small-intestinal-epithelium?filename=regional_cell_sampling_UMIcounts.txt.gz). For each region, we extracted cells annotated as TA, EP and enterocyte. Of these, Mki67-positive cells were defined as crypt cells whereas Mki67-negative cells were defined as villus enterocytes. For each cell, expression data was normalized by the summed cellular UMIs and zonation profiles were reconstructed as described above.

### Clustering and gene ontology enrichment

Gene ontology (GO) terms were obtained from Ensembl (GRCm38 release 90). All GO annotations that contained more than three genes with highly expressed zonated enterocyte genes (UMI fraction above 10^−4^, 2118 genes) were chosen for enrichment analysis. The expression profiles along the villus-axis of these genes were normalized to their maximum expression. The normalized profiles were partitioned in five mutually exclusive clusters with k-Means clustering using MATLAB by using correlation as distance measure (Supp. Table 5). The significance of enrichment of the selected GO terms in each of these five clusters was assessed with the hypergeometric test. Storey’s method was used to compute q-values (Supp. Table 6).

## Sequencing data availability

The generated sequencing data have been deposited in the Genbank GEO database (http://www.ncbi.nlm.nih.gov/geo/) under accession code GSE109413.

## Acknowledgments

A.E.M. is supported by the Swiss National Science Foundation (grant 158999) and the EMBO Long-Term Fellowship program (ALTF 306-2016). S.I. is supported by the Henry Chanoch Krenter Institute for Biomedical Imaging and Genomics, The Leir Charitable Foundations, Richard Jakubskind Laboratory of Systems Biology, Cymerman-Jakubskind Prize, The Lord Sieff of Brimpton Memorial Fund, the I-CORE program of the Planning and Budgeting Committee and the Israel Science Foundation (grants 1902/ 12 and 1796/12), the Israel Science Foundation grant No. 1486/16, the EMBO Young Investigator Program and the European Research Council under the European Union’s Seventh Framework Programme (FP7/2007-2013)/ERC grant agreement number 335122, the Bert L. and N. Kuggie Vallee Foundation and the Howard Hughes Medical Institute (HHMI) international research scholar award. S.I. is the incumbent of the Philip Harris and Gerald Ronson Career Development Chair.

## Author contributions

A.E.M. and S.I. conceived the study. A.E.M. designed and performed most of the experiments. A.E.M., Y.H., E.E.M. and S.B.M. performed single molecule FISH experiments. A.E.M. performed laser capture microdissection experiments. A.E.M. and K.B.H. performed RNA sequencing experiments. S.I. and A.E.M. performed data analysis. S.I. and A.E.M wrote the manuscript. S.I. supervised the study. All authors discussed the results and commented on the manuscript.

## Online Supplementary Material

Supplementary Legends and Supplementary Figures 1-10

Supplementary Tables 1-7

